# Effects of Postmortem Intervals on Quantitative MRI in Unfixed and Fixed Swine Brain: Implications for Ex Vivo MRI Applications

**DOI:** 10.1101/2025.10.08.681281

**Authors:** Lixian Wang, Naoya Oishi, Shin-ichi Urayama, Takashi Hanakawa

## Abstract

**Purpose:** The postmortem interval (PMI) alters tissue properties that shape quantitative MRI (qMRI) signals. We systematically investigated the effects of PMI on relaxation times, depending on tissue characteristics, in both unfixed and fixed pig brains.

**Methods:** Twelve pig brains (n = 12) were scanned in both unfixed and fixed states at three PMI windows (≈12, 24, 48 h). Quantitative T1, T2, and T2* maps were acquired with identical protocols at controlled room temperature (∼24 °C). We assessed the PMI effects on relaxation times in the gray matter (GM) and the white matter (WM) at both group and sample levels, while fixation-induced effects and inter-sample variability were evaluated using pairwise rank correlation and coefficient of variation (CV) visualization.

**Results:** In unfixed tissues, T1 significantly differed among PMI groups in GM (p = 0.0141) and WM (p = 0.0315) in the early window (≤ 24 h). In the same window, PMI–T1 correlations were observed in both GM (r = 0.921, p = 0.007) and WM (r = 0.876, p = 0.013). T2 showed no group differences but exhibited an inverse correlation with PMI in WM (r = −0.849, p = 0.015). No significant PMI–T2* relationships were detected (all p > 0.05). At later PMI (20–50 h), PMI–qMRI correlations diminished. Fixation processes altered all qMRI parameters. Notably, the PMI effects on T1 in the unfixed brains were preserved even after fixation at an ordinal level, although fixation introduced substantial inter-sample variability.

**Conclusions:** PMI exerts the most robust effects on T1. Fixation has a significant impact on qMRI values, mitigating the apparent PMI effect and increasing the inter-sample variability. However, it is still possible to retrieve the PMI effects from fixed brain at the level of rank order, providing practical guidance for further ex vivo qMRI studies on PMI.

## Introduction

*Ex vivo* magnetic resonance imaging (MRI) has become an increasingly powerful tool in basic medicine and science. Extended scan durations enable *ex vivo* MRI to achieve an extraordinarily high spatial resolution because there are no issues with physiological motions (e.g., cardiac and respiratory cycles), restrictions on specific absorption rate (SAR), or the need for living organisms to remain still. *U*ltrahigh-resolution *ex vivo* MRI, achieved through improved signal-to-noise ratio (SNR), leads to the development of notable datasets, including ultrahigh-resolution, multimodal MRI data from both human and animal brains ^1–3^. Detailed investigations of brain microstructure and cross-modal correlations combined with histological analyses are now feasible using *ex vivo* MRI ^4–7^. Quantitative MRI (qMRI), in particular, benefits greatly from the *ex vivo* setting, providing reproducible metrics (e.g., T1, T2, and T2*) that are minimally influenced by physiological variability, thereby making it an ideal approach for radiology–histology/pathology correlation studies ^8–10^.

However, qMRI values derived from *ex vivo* MRI are highly sensitive to both technical and biological variables, particularly tissue fixation and the postmortem interval (PMI) ^11,12^. The effects of formalin fixation on MRI parameters have been addressed, including influences related to nuclear magnetic resonance (NMR) principles, temperature, fixative concentration, and even differences in manufacturers producing fixative solutions ^13–15^. In contrast, the impact of PMI on *ex vivo* qMRI remains understudied. In *ex vivo* MRI, PMI is defined as the interval from the death of an organism to the start of imaging for unfixed specimens, or to the initiation of fixation for fixed specimens (either immersion or perfusion with fixative). Although rodent studies have demonstrated that PMI can significantly affect relaxation and diffusion properties ^16^, findings in human brain tissue indicate minimal effects ^17^. These inconsistencies underscore the need for systematic evaluation. Most previous *ex vivo* MRI studies on PMI have relied on small excised samples of fixed tissue (e.g., cortical slices), which may not fully capture the spatial heterogeneity of MRI signals throughout the entire brain ^16^. Moreover, few studies have conducted direct comparisons between unfixed and fixed tissues derived from the same postmortem brain. This gap is particularly significant, given that fixation often commences several hours or even days after death due to practical and ethical constraints—especially in human autopsy cases. Understanding how MRI signals change as a function of PMI differentially between the fixed and unfixed conditions is essential for the accurate interpretation of *ex vivo* imaging data. Furthermore, these changes may differentially affect the gray and white matter structures, thereby complicating quantitative assessments across the brain.

In this study, we investigated the impact of PMI on qMRI parameters (T1, T2, and T2*) using whole pig brains as a model system. We chose domesticated swine (*Sus scrofa*) due to the anatomical and structural similarities between their brains and the human brain ^17,18^. To isolate the effects of PMI, all other variables—such as fixation buffer, fixation duration, degassing, and phosphate-buffered saline (PBS) immersion procedures—were held constant across all samples. We performed imaging of unfixed brains at three PMIs: approximately 12, 24, and 48 hours. We evaluated how PMI affected key qMRI metrics in both unfixed and fixed tissue conditions, using an identical MRI protocol. Our central hypothesis was that PMI would significantly alter qMRI values in unfixed brain tissue. We expected that fixation might partially mask these effects, with different effects on the gray matter (GM) and white matter (WM), but also assumed that the PMI effects could still be observable after fixation. Knowledge from this research should enhance the interpretation of human *ex vivo* qMRI data, particularly in settings where precise control over PMI is not feasible. Ultimately, we aim to provide insights into *ex vivo* human brain research, thereby enhancing the translational value of qMRI datasets from *ex vivo* to *in vivo* applications.

## 2. Materials and Methods

### 2.1 Specimen Preparation and Experimental Design

We used twelve brains extracted from domesticated swine (*Sus scrofa*) in this study. All specimens were procured from a local wholesale market (Kyoto Meat Market). No animals were slaughtered for this study. Animal handling and tissue acquisition were conducted in compliance with local regulations (Japan Slaughterhouse Act, Act No. 114 of 1953). Four brains were assigned to each of the three experimental groups defined by the target PMI: the 12 hours, 24 hours, and 48 hours PMI groups. The details of the groups and experimental timelines are listed in Table 1.

**Table 1.** Grouping information, including PMI fixation duration and PBS immersion duration.

| Groups | Samples | PMI<br>(unfixed,<br>T1&T2*<br>scan, h) | PMI<br>(unfixed,<br>T2 scan, h) | PMI (fixed<br>scan, h) | Fixation<br>(days) | PBS<br>immersion<br>(days) |
| --- | --- | --- | --- | --- | --- | --- |
| PMI<br>12 h | ① | 9 | 10 | 11 | 30 | 14 |
|  | ② | 7.5 | 8.5 | 9.5 | 30 |  |
|  | ③ | 11 | 12 | 13 | 29 |  |
|  | ④ | 9 | 10 | 11 | 30 |  |
|  | mean±SD | 9.13±1.24 | 10.13±1.24 | 11.13±1.24 | 29.75±0.43 |  |
| PMI<br>24 h | ① | 22.5 | 23.5 | 24.5 | 30 |  |
|  | ② | 20 | 21 | 22 | 29 |  |
|  | ③ | 23.5 | 24.5 | 25.5 | 29 |  |
|  | ④ | 22 | 23 | 24 | 30 |  |
|  | mean±SD | 22±1.27 | 23±1.27 | 24±1.27 | 29.5±0.50 |  |
| PMI<br>48 h | ① | 44.5 | 45.5 | 47 | 29 |  |
|  | ② | 48 | 49 | 50 | 29 |  |
|  | ③ | 44 | 45 | 46.3 | 29 |  |
|  | ④ | 47.5 | 48.5 | 49.7 | 29 |  |
|  | mean±SD | 46±1.77 | 47±1.77 | 48.25±1.62 | 29 |  |

PMI, postmortem interval; PBS, phosphate-buffered saline; SD, standard deviation

### 2.2 Experimental Procedure

We extracted the cerebrum from the skull, followed by the removal of the dura mater and arachnoid membrane, to mitigate the risk of air bubbles being trapped in the subarachnoid space after fluid immersion. The average mass of a typical pig brain is approximately 130 g, roughly one-tenth that of the adult human brain ^18^. The mass of the cerebrum, excluding the meninges, was approximately 110 g. All samples were extracted within 12 hours after the death and subsequently subjected to a degassing procedure.

We performed degassing with a vacuum pump and chamber. For degassing of fixed specimens, a near-vacuum condition is widely employed because an extremely low-pressure vacuum enhances gas removal. Degassing is often considered complete when no apparent bubbling is observed at ∼10 mbar. However, the near-vacuum degassing procedure damages brain tissue ^19^, and our unfixed specimens are even more susceptible to damage ^20^. To prevent damage to delicate, unfixed brain tissue, we followed a gentler degassing protocol at ∼250 mbar ^21^. Specifically, specimens were immersed in PBS (Nacalai Tesque, Kyoto, Japan) and degassed at ∼200 mbar for one hour by vacuum pump (DTC-21, Ulvac, Inc., Kanagawa, Japan) and chamber.

To ensure stable positioning during imaging and to preserve the shape of the unfixed brain specimens, we designed and fabricated a custom-made, three-dimensional (3D)-printed brain holder shaped to resemble a pig skull (Figures 1A and B) by a 3D printer (Agilista 3000, Keyence, Osaka, Japan). We designed and created a lid to accommodate probes for real-time temperature monitoring. This brain holder enabled secure placement of the specimen within a cylindrical container filled with PBS, which was then inserted into an MRI coil (Figure 1C). The temperature of the fluid surrounding the specimen was continuously monitored using fiber optic probes (FISO, Quebec, Canada; model: Evolution EVO-SD-4; serial number: TT8237A) (Figure 1D) in approximately 50% of the MRI sessions.

**Fig. 1.**
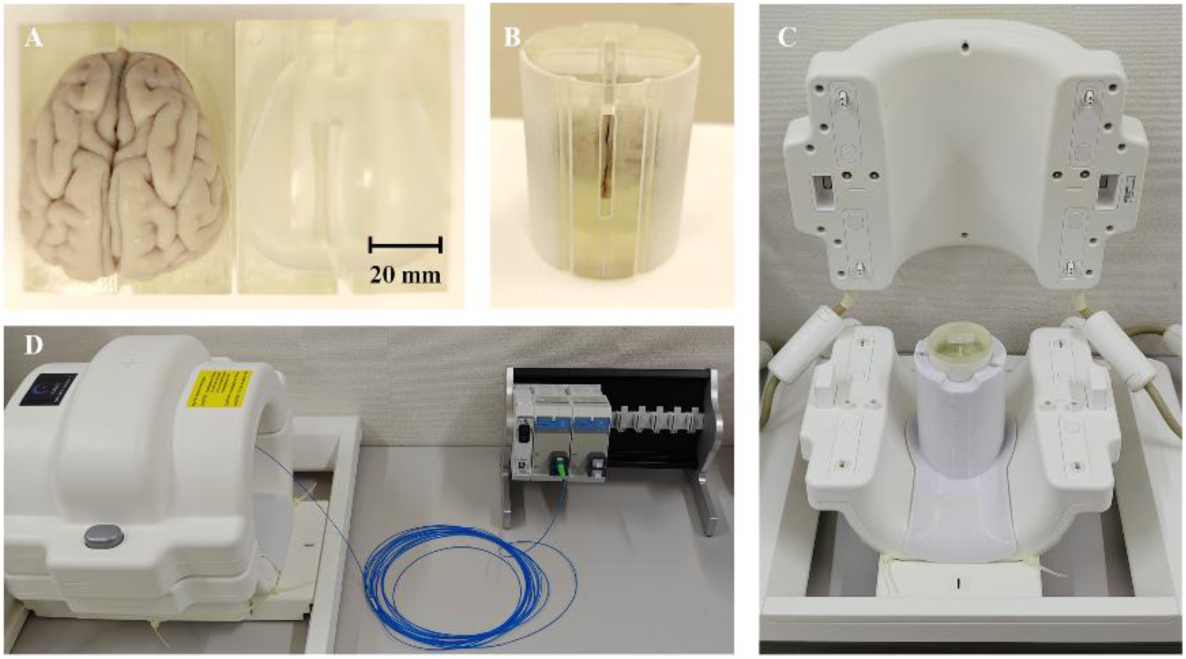
Materials Used for MRI Scanning. (A) Pig brain and 3D-printed holder before assembly. (B) Pig brain mounted within the 3D-printed holder. (C) Knee coil, insertable coil, and the assembled sample. (D) The temperature monitoring system.

The containers were sealed to prevent the ingress of additional air. Despite the degassing procedure, small air bubbles often remained during the initial scan conducted at ∼12h PMI. After the seal, the unfixed specimens were stored in the MRI room to minimize temperature fluctuations until their designated PMI time points (∼12, 24, or 48 hours). In these subsequent scans, most air bubbles had dissolved into the PBS. All procedures were conducted at the MRI room temperature (approximately 24 °C).

Immediately after the scanning of the unfixed samples, the brains were fixed in 10% neutral buffered formalin (NBF; Nacalai Tesque, Inc., Kyoto, Japan). While immersing the sample in the fixative, the 3D-printed holder was kept attached for an additional two days to maintain the sample’s shape. After approximately 30 days of fixation, the samples were rinsed with and immersed in PBS to eliminate residual fixative. The brains were immersed in PBS for two weeks, with the PBS replaced after two days to ensure thorough dilution of the fixative. Following washing, the fixed samples were returned to the 3D-printed holder and scanned again using the identical MRI protocols.

Figure 2 summarizes the experimental workflow and timeline. Only the interval between degassing and the unfixed scan differed, as determined by the target PMI. We conducted a total of four scanning sessions, two before and two after fixation: (i) T1&T2* scanning of unfixed specimens with multi-echo gradient echo (MEGRE) sequence; (ii) T2 scanning of unfixed brain with turbo spin-echo (TSE) sequence; (iii) T1&T2* scanning of fixed brains with MEGRE; and (iv) T2 scanning of the fixed brains with TSE. For the scanning of the unfixed brain, actual PMI was recorded as the interval from death to the start of the T1&T2* or T2 scanning. For the fixed brain, actual PMI was recorded as the interval from death to the initiation of formalin fixation, assuming that the PMI process is ceased at fixation. The details of PMI definitions and specimen conditions, such as fixation duration and PBS immersion duration, are provided in Table 1.

**Fig. 2.**
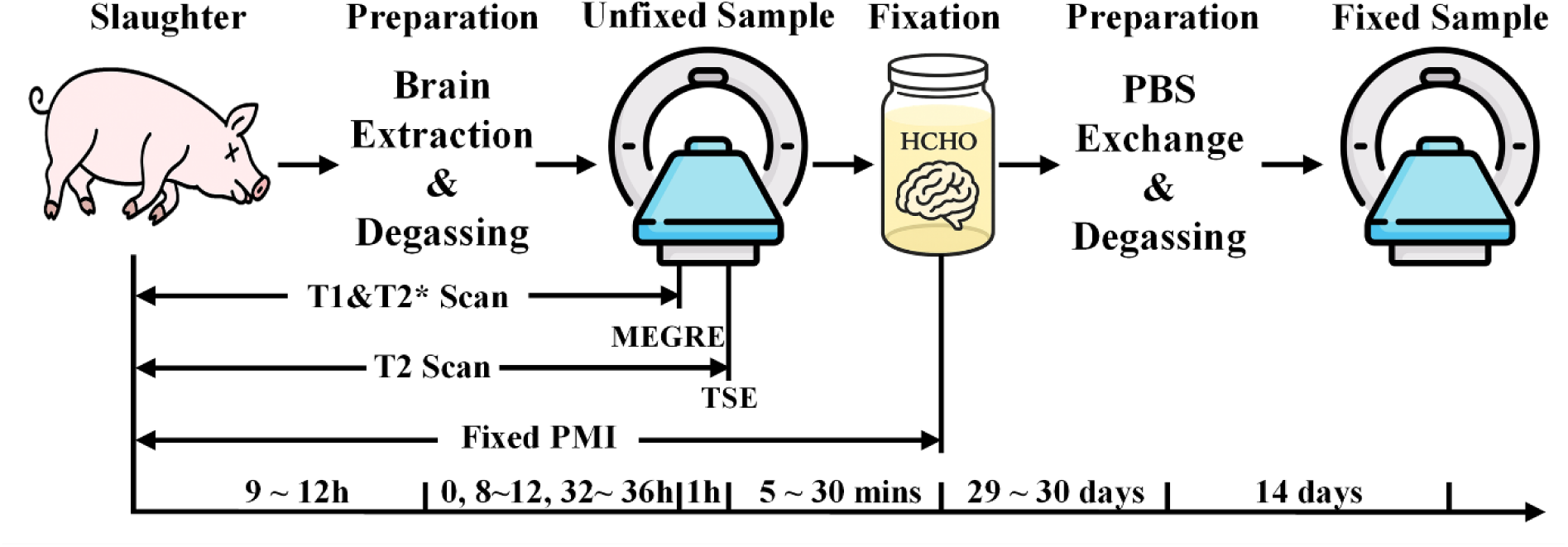
Experimental workflow and timeline, including the definition of PMI PBS, phosphate-buffered saline; MEGRE, multi-echo gradient echo; TSE, turbo spin echo; PMI, postmortem interval

### 2.3 MRI Acquisition

MRI experiments were conducted using a whole-body 7T scanner (MAGNETOM 7T, Siemens Healthineers, Erlangen, Germany) equipped with a single-channel transmit and 28-channel receiver coil designed for knee scanning (Quality Electrodynamics, Mayfield Village, OH, USA) ^22^ (Figures 1C & D). To enhance SNR, an insertable, inductively coupled volumetric coil with an inner diameter of 64 mm was employed (Figure 1C) ^23^.

For T1 and T2* mapping, a 3D MEGRE sequence was adopted with five flip angles, each with eight echo times. B1 maps were acquired using the SAturation prepared with 2 RApid Gradient Echoes (SA2RAGE, Siemens prototype sequence) ^24^. For T2 mapping, a TSE sequence was acquired with four echo times. High-resolution 3D T1-weighted (T1w) structural images were acquired using the Magnetization-Prepared 2 Rapid Acquisition Gradient-Echoes (MP2RAGE, Siemens prototype sequence). The sequence parameters are listed in Table 2. All MRI sequences were performed on both unfixed and fixed samples.

**Table 2.** MRI sequences and parameters.

| Target | Sequence | TA | TR | TE | FA | Resolution | Other |
| --- | --- | --- | --- | --- | --- | --- | --- |
| **T<sub>1</sub> & T2\***<br>**Mapping** | 3D<br>MEGRE | 12 mins<br>×5 | 46<br>ms | 3.4, 6, 8.6,<br>11.3, 13.9,<br>16.5, 19.1,<br>21.7 ms | 7°, 17°,<br>27°, 37°,<br>47° | Isotropic<br>0.4 mm | BW: 500Hz |
| **T<sub>2</sub>**<br>**Mapping** | 3D TSE | 17 mins<br>×4 | 500<br>ms | 11, 34, 56,<br>79 ms |  |  |  |
| **Structure** | MP2RAGE | 13 mins | 6000<br>ms | 3.3 ms | FA1/FA2:<br>4°/5° |  | TI1/TI2:<br>800/2700ms |
| **B<sub>1</sub>**<br>**Mapping** | SA2RAGE | 2 mins | 2400<br>ms | 0.88 ms | FA1/FA2:<br>4°/10° | Isotropic<br>1.6 mm | TI1/TI2:<br>50/1800ms |

TSE, turbo spin echo; TA, acquisition time; TR, repetition time; TE, echo time; FA, flip angle;
MEGRE, multi-echo gradient echo; BW, bandwidth; TI, inversion time

### 2.4 Data Processing

Given the necessity of fine PMI control for unfixed specimens, we performed simultaneous quantification of T1 and T2* values by MEGRE sequences. T1 mapping employed the variable flip angle (VFA) approach, using the shortest TE images from the variable FAs MEGRE data ^25^. B1map from SA2RAGE sequence to correct flip angle’s inaccuracies arising from B1+ inhomogeneity in T1 mapping ^26^. Since the B1 map of one sample was unreliable, T1 maps were generated for 11 fixed specimens. T2* maps were obtained from the lowest FA MEGRE data using mono-exponential fitting^27^. T2 maps were estimated from the TSE data by fitting magnitude images using the same procedure as for T2* mapping. Quantitative T1, T2, and T2* maps were computed using the qMRLab toolbox ^28^ in MATLAB (R2024a; MathWorks, Natick, MA, USA).

Image registration was performed using the Advanced Normalization Tools (ANTs) ^29^. Rigid-body registration was applied to align all images to the first echo of the first flip angle from the 3D MEGRE sequence, to minimize potential bias in subsequent quantitative map calculations. This reference image was subsequently used for registration to the 3D T1-weighted (T1w) image ^30^.

Brain extraction was performed using the Brain Extraction Tool (BET) ^31^ from the FMRIB Software Library (FSL, version 6.0.7.11) ^32^. The brain mask, generated from the final T1w image and the second inversion recovery (IR) pulse image of the MP2RAGE, was applied to the quantitative maps and subsequent processing steps.

Following the brain image extraction, N4ITK bias field correction was applied to all T1w images ^33^. The bias-corrected images were then used to construct a study-specific template using ANTs ^34^. These bias-corrected T1w images were also used for tissue segmentation using FMRIB’s Automated Segmentation Tool (FAST), which generated tissue probability maps ^35^. These maps served as priors to improve segmentation accuracy for each individual sample. After the segmentation, voxels labelled as GM or WM with a probability of 100% were used to construct masks for tissue-level analyses of group differences and correlations with PMI. After the analyses were performed in individual space, the results were spatially normalized to the template using ANTs for improved visualization. An overview of the data processing pipeline is shown in Figure 3.

**Fig. 3.**
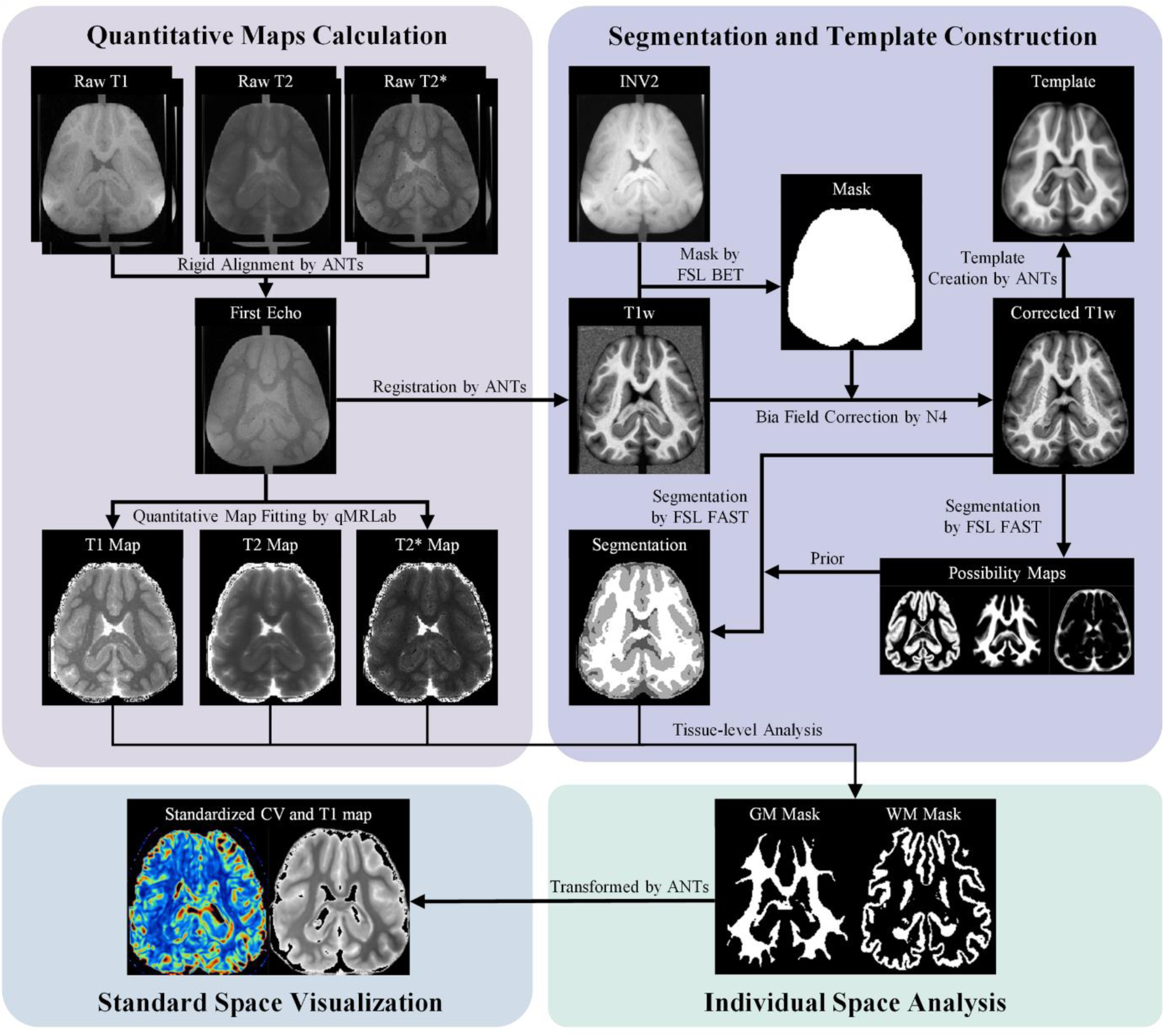
Workflow for Quantitative MRI Processing INV2, the second inversion recovery pulse image of the MP2RAGE; GM, gray matter; WM, white matter; T1w, T1-weighted

### 2.5 Statistical Analysis

Following image processing, group-level comparisons across PMI groups were conducted using analysis of variance (ANOVA) on T1, T2, and T2* values in GM and WM. When normality and homoscedasticity (Brown–Forsythe *p* ≥ 0.05) were satisfied, one-way ANOVA was applied; when variances were unequal (Brown–Forsythe *p* < 0.05), Welch’s ANOVA was used, and Welch-adjusted F- and p-values were reported. Following ANOVA, *post hoc* pairwise comparisons were performed to identify specific group differences. When homoscedasticity was met, Tukey’s HSD was applied; when variances were unequal, the Games–Howell test was used. Statistical significance was set at α = 0.05. All statistical analyses and visualizations were performed in GraphPad Prism (version 10.5.0; GraphPad Software, Inc., San Diego, CA, USA).

To characterize the influence of PMI, relationships between PMI and quantitative relaxation times (T1, T2, and T2*) were assessed in both unfixed and fixed specimens. The PMIs for the T1, T2, and T2* acquisitions and for formalin fixation varied by several hours across specimens. This variability may be biologically meaningful for interpreting postmortem changes (e.g., neuropil vacuolization increases to ∼24 h PMI and decreases by ∼72 h ^36^), particularly at early postmortem stages. Therefore, PMIs were further treated not only as a categorical variable in statistical analyses but also as a continuous parameter. To assess consistency with prior work reporting postmortem effects within ∼24 h PMI ^16^, separate analyses were performed for specimens with PMI ≤ 24 h and those between 20 and 50 h. Linear regression was performed using the least-squares method, and 95% confidence intervals (CIs), the correlation coefficient (r), and corresponding p-values were computed. To correct for multiple comparisons, the false discovery rate was controlled at q = 0.05 using the Benjamini– Hochberg procedure ^37^. We utilized the segmented GM and WM masks to retrieve tissue-level (GM and WM) qMRI associations with PMI.

The measurements before and after fixation were visualized using paired violin plots with overlaid points and within-specimen linking lines, stratified by PMI group (12 h, 24 h, and 48 h), tissue class (GM and WM), and relaxation parameter (T1, T2, and T2*). We calculated the Spearman rank correlation coefficient between unfixed and fixed measurements. Given the very small sample size per PMI group, we reported ρ values only in each PMI group, treating these as descriptive indicators of monotonic association. To further assess whether the rank order of quantitative values was preserved after fixation, we computed Spearman’s rank correlation using a PMI-wise rank pooling approach. The qMRI data were first ranked within each PMI group. The rank order was pooled across all groups, yielding 12 paired ranks fed into Spearman’s correlation analysis. Spearman’s ρ and the two-tailed p-value were reported.

For visualizing the effects of PMI and fixation on the T1 relaxation times, coefficients of variation (CVs) were computed for each group as the SD divided by the mean T1, and are reported as descriptive summaries. Mean T1 and CVs were presented as descriptive summaries.

## 3. Results

Given the temperature sensitivity of qMRI parameters, all procedures were conducted at near-room temperature (approximately 24 °C). Temperature was recorded as frequently as feasible by monitoring the temperature of the surrounding PBS. The initial temperature for the T1 and T2* protocol was 24.57 ± 1.24 °C (mean ± SD). Following T1 measurements, temperature increased by 1.04 ± 0.59 °C; following T2* measurements, the increase was 0.02 ± 0.01 °C. The initial temperature for the T2 protocol was 25.64 ± 0.86 °C, and after T2 measurements, the temperature increased by 1.44 ± 0.52 °C. According to prior reports, the magnitude of these fluctuations is unlikely to affect the qMRI measurements ^11^. Therefore, temperature was not included as a covariate in subsequent analyses. We evaluated the effects of PMI on relaxation times. In the unfixed samples, statistically significant correlations were identified in several tests. Group-level PMI effects on relaxation times were analyzed using one-way ANOVA or Welch ANOVA tests (Table 3) and summarized through pairwise patterns with *post hoc* tests from the bar plots (Figure 4). T1 differed across PMI groups in GM (PMI 12 h = 1611 ± 20 ms; PMI 24 h = 1689 ± 28 ms; PMI 48 h = 1676 ± 42 ms; p = 0.0141) and WM (1251 ± 12, 1309 ± 29, 1319 ± 52 ms; p = 0.0315). *Post hoc* comparisons (Figure 4) indicate higher T1 at longer PMI (12 h < 24/48 h; 24 h ≈ 48 h). No differences in T2 were detected for GM (p = 0.823) or WM (p = 0.310). T2* showed a group effect in GM (50.78 ± 0.48, 51.96 ± 0.82, 53.47 ± 1.76 ms; p = 0.0474), while *post hoc* contrasts did not reach significance. T2* showed no group effect in WM (p = 0.499).

**Fig. 4.**
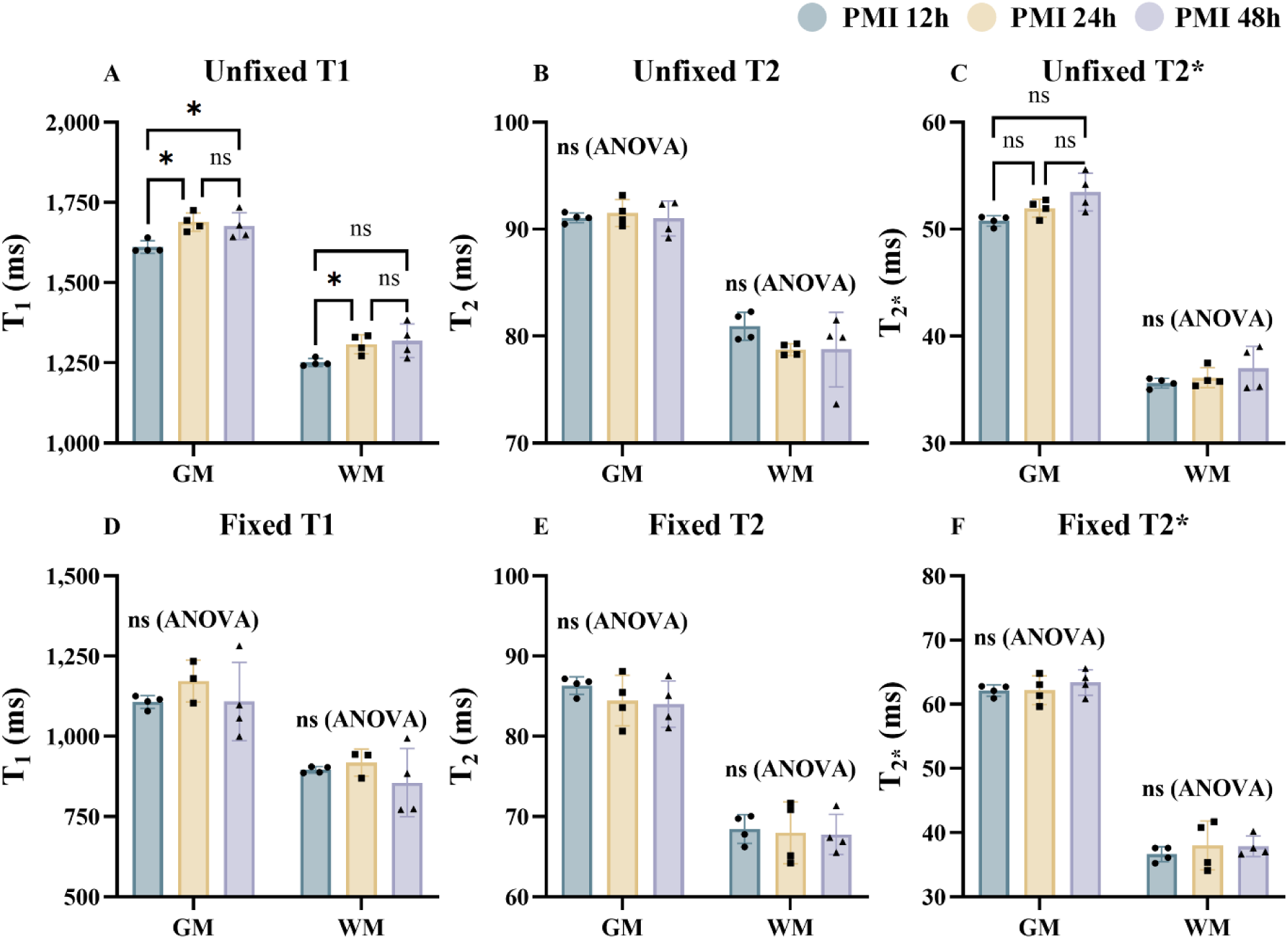
Group-level difference in qMRI parameters across PMI groups in unfixed and fixed brain samples PMI, postmortem interval; GM, gray matter; WM, white matter; *, significant difference in *post hoc* test (p < 0.05); ns, not significant in *post hoc* test; ns (ANOVA), not significant in analysis of variance

**Fig. 5.**
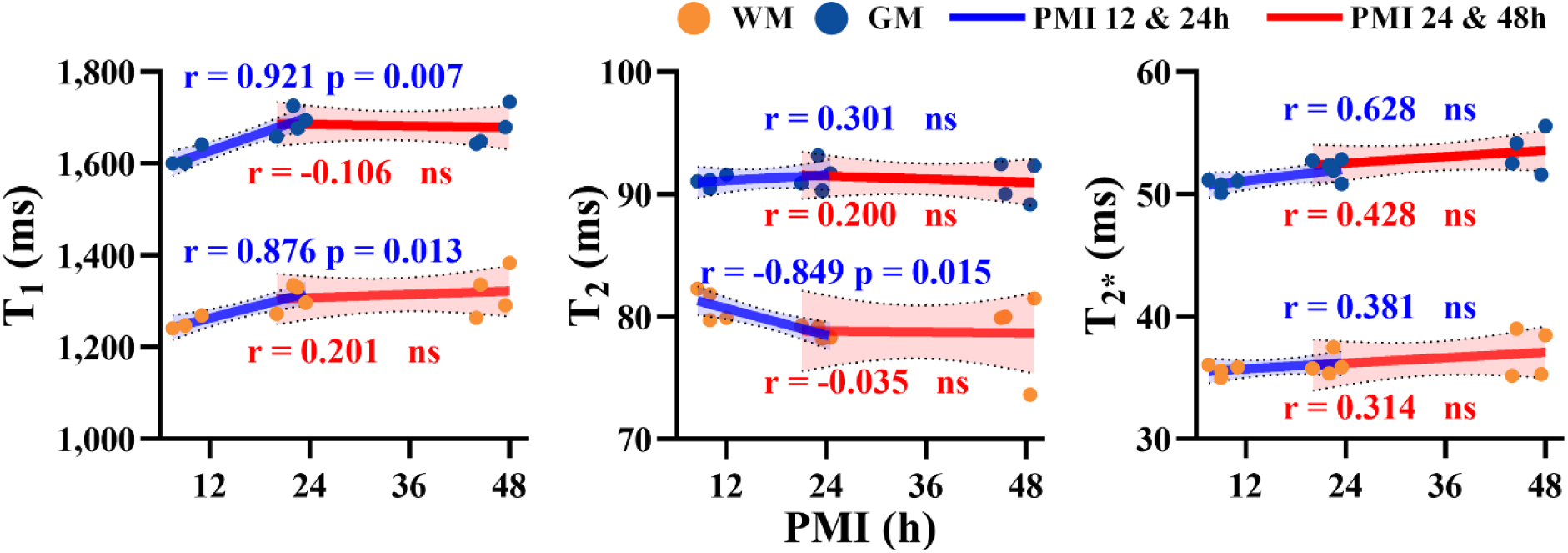
Tissue-level correlations between T1, T2, and T2* relaxation times and PMI in unfixed brain specimens WM, white matter; GM, gray matter; PMI, postmortem interval; ns, not significant

**Fig. 6.**
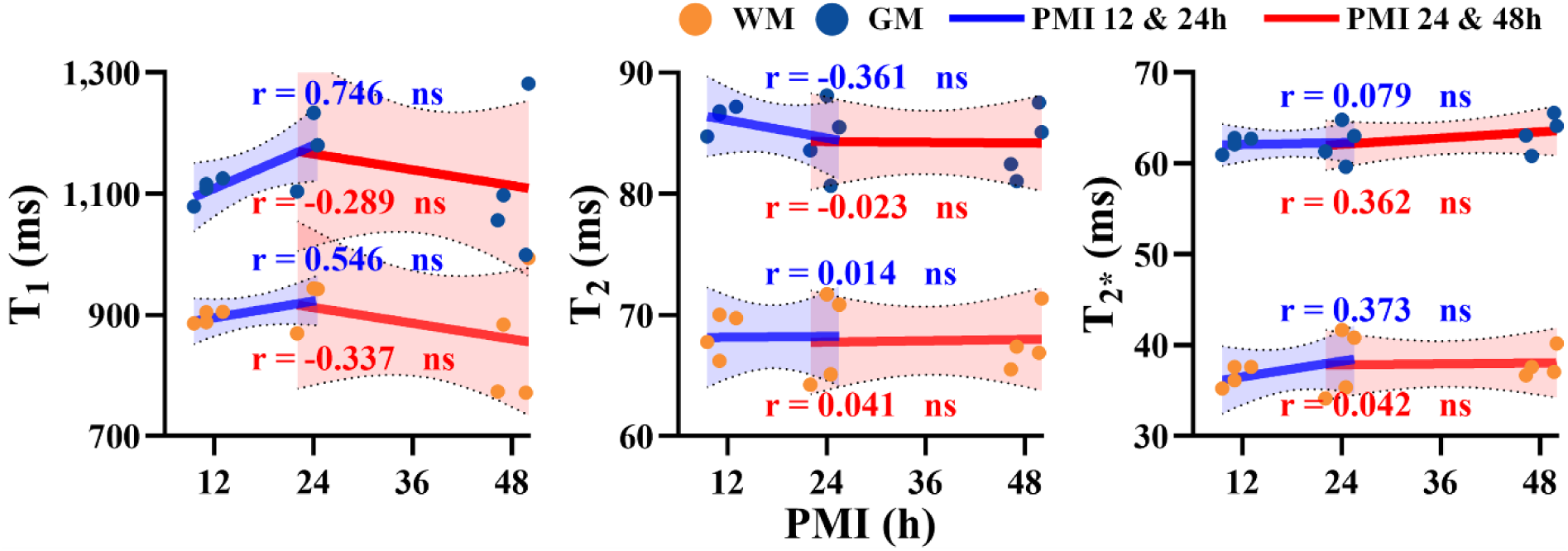
Tissue-level correlations between T1, T2, and T2* relaxation times and PMI in fixed brain specimens WM, white matter; GM, gray matter; PMI, postmortem interval; ns, not significant

**Table 3.**
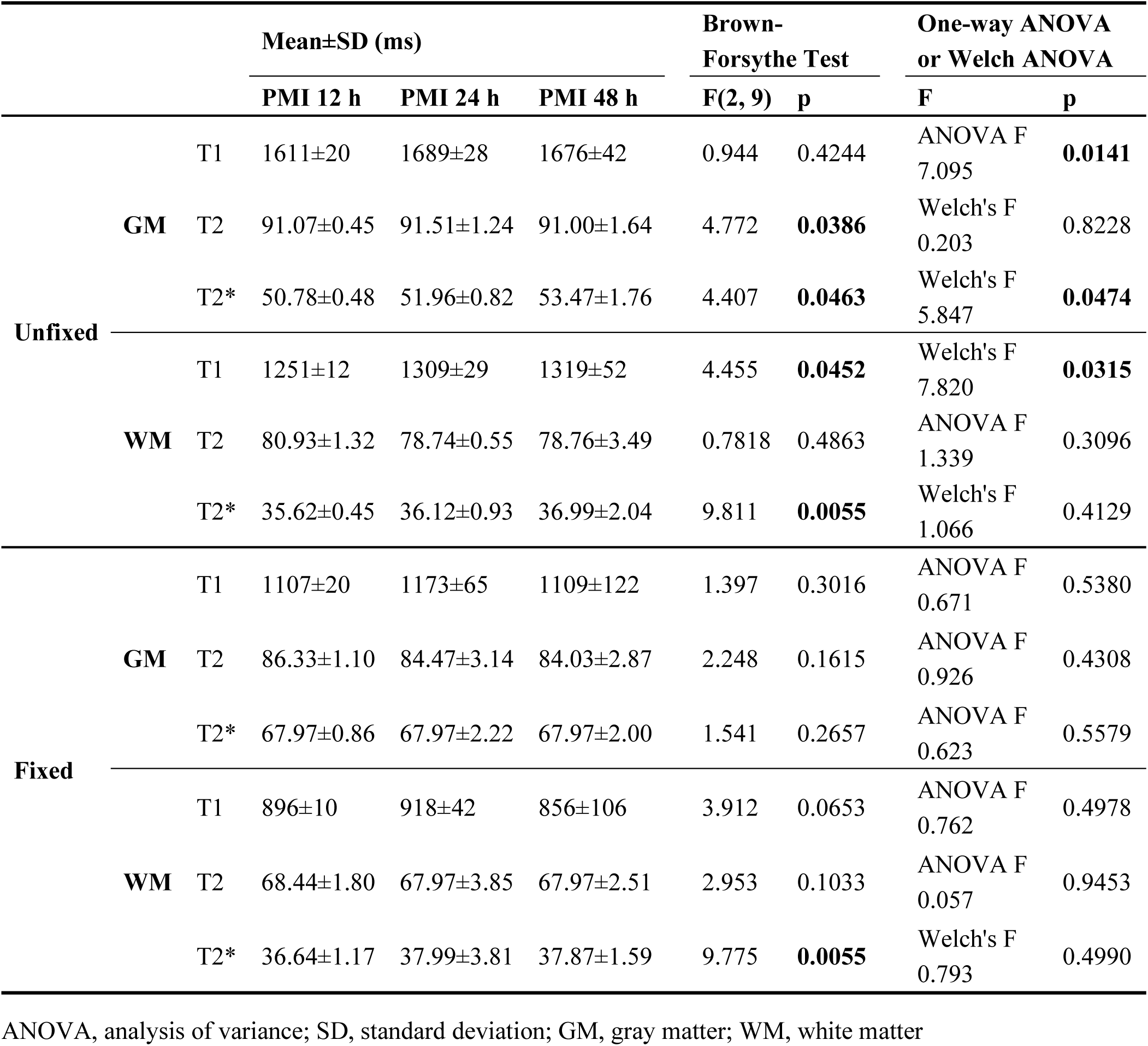
Summary statistics and results from ANOVA on quantitative MRI parameters from unfixed and fixed swine brain samples.

Correlation analyses were conducted to investigate the more nuanced effects of PMI on relaxation times, taking into account subtle differences in specimen preparation. Within the first ∼24 hours postmortem, T1 relaxation time showed a strongly positive association with PMI in the GM (r = 0.921, p = 0.007) and a weaker yet positive association in the WM (r = 0.876, p = 0.013). T2 relaxation time exhibited a significantly negative association with PMI in the WM (r = -0.849, p = 0.015), but not in the GM. No significant association was observed between PMI and T2* relaxation time in either GM or WM. In the unfixed samples with PMI between 20 and 50 hours, tissue-level analyses revealed no significant associations between PMI and relaxation times (T1, T2, or T2*).

After formalin fixation, T1 and T2 relaxation times significantly decreased, while T2* increased (Table 3 and Figure 7), consistent with previous report ^38^. Notably, group-level differences in the PMI effects almost disappeared, and the correlations between relaxation times and PMI observed in the <24h postmortem unfixed specimens also disappeared after the fixation procedure (Figure 8 A and B). The loss of the statistically significant PMI effects on relaxation times appeared to result from, at least in part, a large data variance after fixation compared with the one before fixation. In the unfixed state, mean T1 increased with PMI in both GM and WM, consistent with the preceding analyses. After fixation, the absolute T1 values became uniformly lower, and the increasing trend with PMI was no longer apparent. The voxel-wise CV (clipped at 15%) across specimens within each PMI group provided information on the data variance (Figure 8C and D). For unfixed brains, variability was lowest at 12 h and modestly higher at 24–48 h (predominantly ≤9%), indicating relatively consistent T1 at shorter PMI. In contrast, fixed brains showed a clear PMI dependence of variability: CV rose from 12 h to 48 h, with widespread elevations (often ≥12%) at 48 h, particularly in cortical GM and deep structures. Taken together, the mean and CV maps indicated that unfixed T1 tended to rise with PMI, fixation shortened T1, and inter-specimen variability was minimized at short PMI but became accentuated at longer PMI, most notably after fixation.

**Fig. 7.**
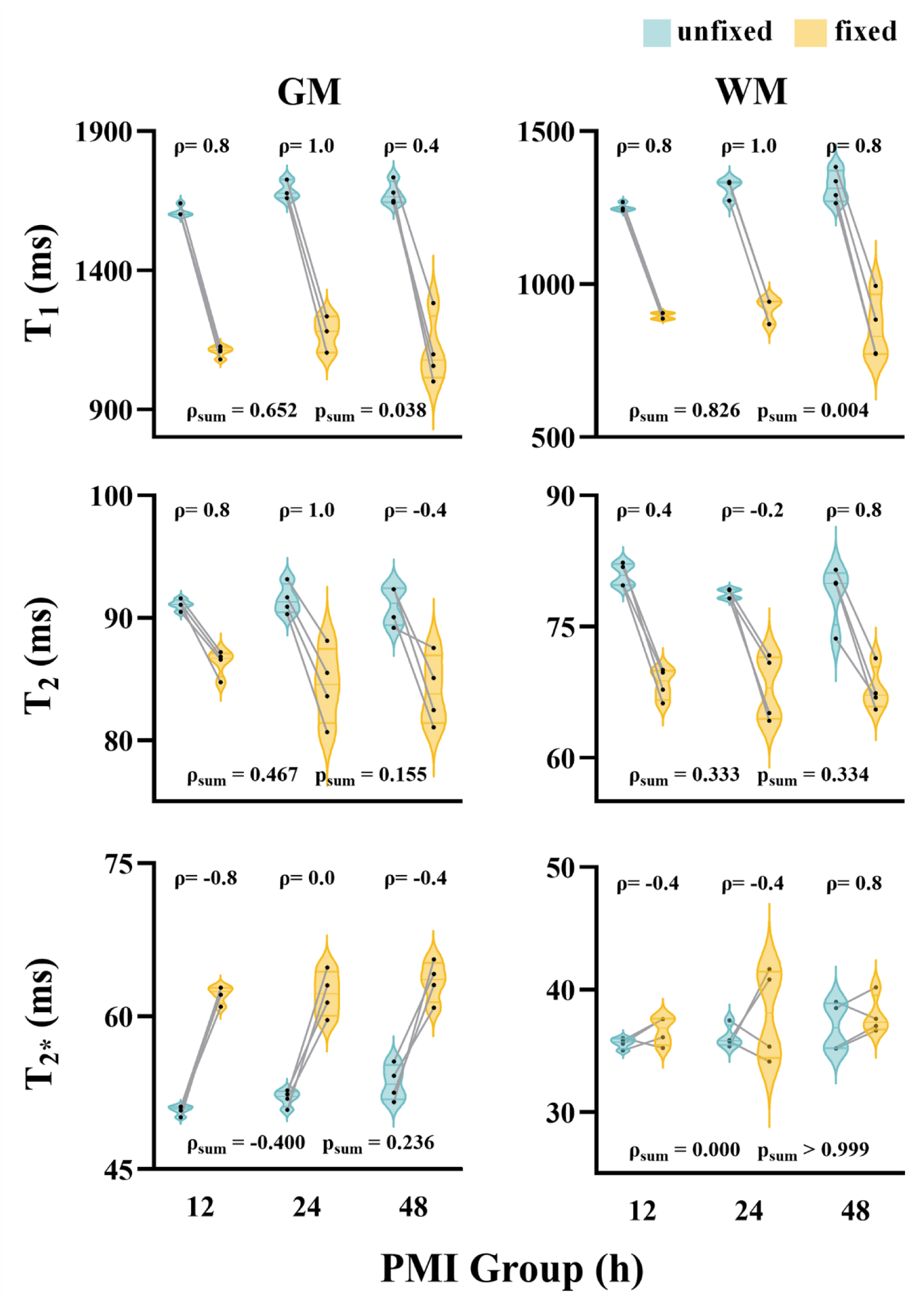
Paired comparisons of quantitative parameters within each PMI group and Spearman’s rank correlation (ρ) between before and after fixation measurements GM, gray matter; WM, white matter; PMI, postmortem interval; ρ, Spearman’s rank correlation coefficient for each PMI group; ρ_sum_, Spearman’s rank correlation coefficient for all samples; p_sum_, significance level for all samples

**Fig. 8.**
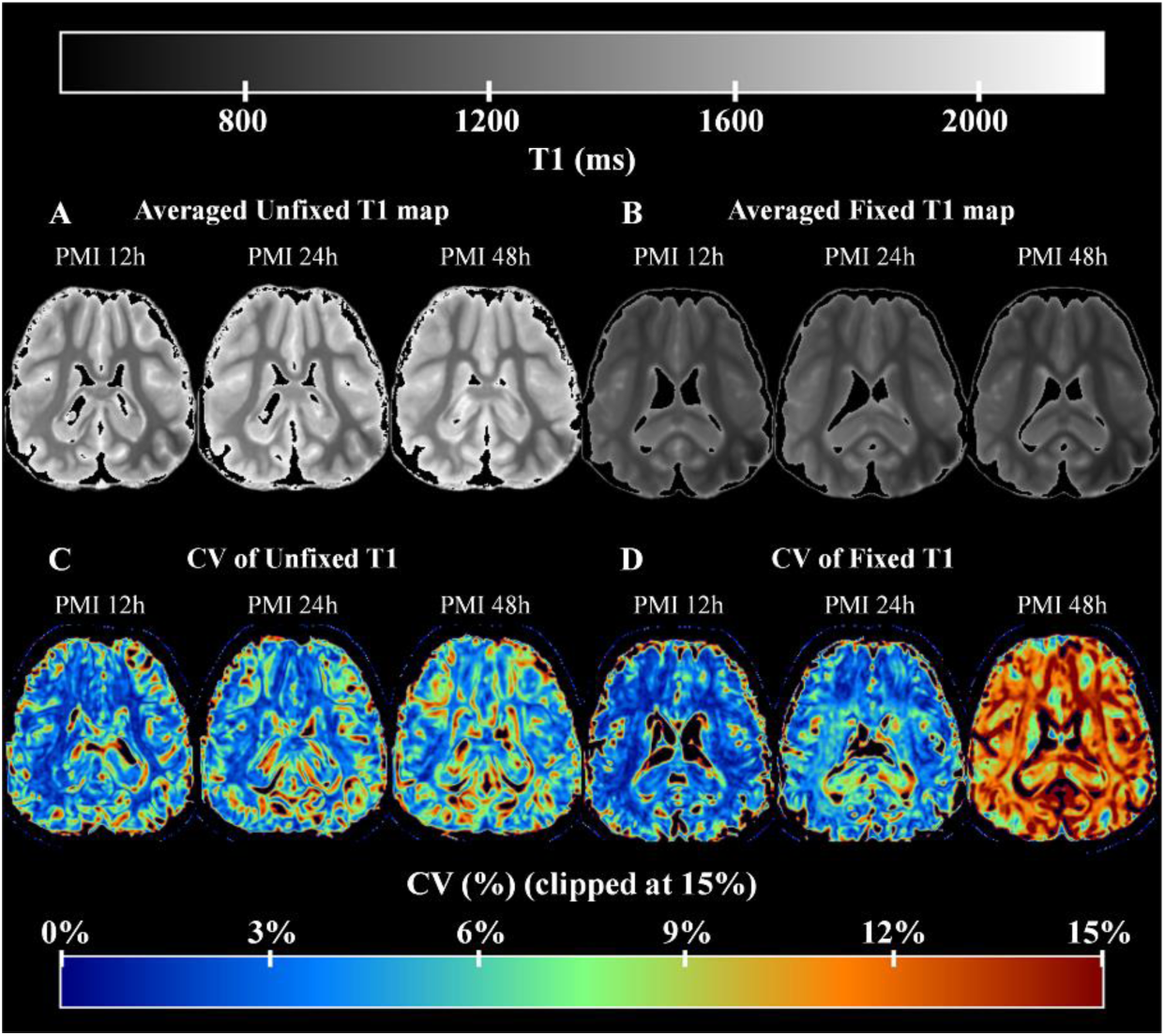
Group-wise T1 comparisons in unfixed and fixed samples: averaged T1 maps (A, B; Unfixed and fixed, both for PMI 12, 24, and 48 h) and voxelwise CV maps (C, D; Unfixed and fixed, both for PMI 12, 24, and 48 h). Mean images share a common display window; CV maps share a common 0– 15% scale (clipped at 15%). PMI, postmortem interval; CV, coefficient of variation

To investigate whether the PMI effects were preserved between fixated and fixated specimens, the Spearman rank correlation coefficient (ρ) was calculated first within each PMI group (Figure 7). Most PMI groups showed high correlations (ρ ≈ 0.8–1.0) for both GM and WM, indicating that the PMI effect was largely preserved after fixation. In contrast, T2 and T2* exhibited more variable and irregular ρ values across PMI and tissue type, with T2* showing the greatest scatter and no consistent pattern. The preservation of PMI effects across both unfixed and fixed specimens was maintained when all PMI groups were combined. The strongest monotonic correspondence between unfixed and fixed measurements was observed for T1, while T2 and particularly T2* displayed more random relationships. For T1, the ordinal preservation of the PMI effect reached statistical significance in both GM (p = 0.038) and WM (p = 0.004). These findings suggest that fixation has a minimal impact on the PMI effect measured by T1 relaxation times from an ordinal perspective, whereas the PMI effect is more fragile when measured using T2 and T2*. Note, however, that fixation increased the variance in all qMRI measures.

## 4. Discussion

We found a substantial effect of PMI on *ex vivo* qMRI. In unfixed specimens, associations between PMI and qMRI parameters were evident and most pronounced within the first 24 h postmortem. These effects varied by quantitative parameter (T1, T2, and T2*) and by the tissue type (GM vs WM). After fixation, these associations were attenuated or disappeared, likely because the effects of formalin on qMRI largely masked the PMI effect ^39^. Nevertheless, PMI influenced measurement consistency (e.g., SD/CV). Therefore, PMI should be reported in postmortem MRI studies wherever possible, controlled where feasible, and explicitly incorporated into analyses.

Our results bridge knowledge gaps in earlier postmortem MRI and histological reports. A prior histology study showed a graded sequence of tissue degradation as a function of PMI—perinuclear vacuolization and neuropil pallor by ∼4–12 h, progressing to retraction artifacts and pyknotic nuclei by ∼24–36 h—consistent with cytotoxic edema and early autolytic change ^16^. Cytotoxic edema produces a progressively more irregular cellular architecture as PMI increases, resulting in a more heterogeneous distribution of water in tissues with prolonged PMIs. This process would differ across specimens due to their intrinsic biological variability, as well as technical differences in procurement, temperature/cooling, and immersion conditions. This inter-sample difference in tissue condition is likely to be pronounced once triggered as PMI increases. Indeed, variances in all relaxation time measures tended to increase from 12 h to 48 h PMIs in this experiment (Table 3). This finding strongly supports the idea that different samples entered the fixation process at varying histopathological stages (e.g., cytotoxic edema to early autolysis). Our finding is consistent with a previous study reporting positive PMI associations in fixed tissue, particularly in GM ^16^. In our data, the PMI-qMRI associations were observed before, but not after, fixation, due to large inter-sample variability after fixation. However, while the rank order of the PMI effects on T1 relaxation was preserved between unfixed and fixed samples (*see* Figure 7), the inter-sample differences were exaggerated by the tissue fixation process. The reason why fixation enhances the inter-sample difference in histopathological stages is plausibly explained by the slow penetration of the fixative (≈0.5–1.0 mm/h) ^39^. In centimeter-scale brains, deep regions remain unfixed for many hours after fixation starts; therefore, substantial tissue volumes continue to undergo postmortem changes. Consequently, the effective PMI—i.e., the time that tissue effectively remains unfixed before local fixation actually begins—becomes longer than the nominal “fixed PMI” and spatially non-uniform, due to the long postmortem processes in the deep brain structures. The various stages of the samples differentially influence the level of protein cross-linking induced by formalin, thereby enhancing differences in the structural architecture that affects the protons’ environment in the tissue, which particularly influences T1 relaxation times. Consistent with our results, this non-uniform extension of effective PMI leads to higher T1 values (particularly in GM) and greater inter-sample variability, which can reshape or mask the associations observed in unfixed tissue. By acquiring data in both unfixed and fixed conditions and explicitly defining PMI windows, this study provides a contextualization of prior findings.

Given the limited systematic work on PMI in *ex vivo* MRI fields, several observations are novel. First, PMI-qMRI associations occurred predominantly within ∼24 h in unfixed tissue, while they were weak or absent thereafter. This timing is consistent with histological reports of rapid early change followed by partial stabilization. Second, sensitivity differed by parameter and tissue: T1 exhibited the most consistent associations for both GM and WM, while T2 showed weaker or region-specific associations in WM, and T2* showed no associations at all. GM demonstrated stronger positive associations than WM, and WM sometimes showed negative associations for T2. To enhance discussions, we summarize the aforementioned changes across parameters and tissues in the present study (see Table 3). This pattern identifies the readouts and tissue class sensitive to PMI. Third, although fixation reduces group-level associations, PMI still influences measurement variability (e.g., lower SD/CV at shorter PMIs) and thus remains relevant post-fixation. Fourth, in some samples, T2* increased after fixation. This counter-directional finding highlights the complexity of *ex vivo* contrasts and points to additional factors—field inhomogeneity and iron-related sources—that can outweigh simple water-content effects in WM ^40,41^. Together, these observations refine when (early), what (T1 > T2 ≫ T2*), and where (GM > WM for positive associations) PMI matters most.

The observed order of sensitivity (T1 > T2 > T2*, Figure 7) is consistent with established tissue physics and biology. Early postmortem water redistribution and loss of membrane integrity are expected; these processes tend to prolong T1 and also modulate T2 relaxation times. Given GM’s higher water content and distinct microstructure, GM tends to exhibit stronger early changes ^36,40,42–44^. Fixation introduces a second, spatially heterogeneous process: cross-linking progresses over time and varies with depth due to the sluggish infiltration of the fixative. Beyond water and membranes, blood-related susceptibility may contribute: shifts in intravascular hemoglobin species (e.g., deoxyhemoglobin, methemoglobin) increase local field inhomogeneity and can shorten T2/T2* ^40,41^. These effects are likely secondary for T1 relative to water-related changes, but they help explain some WM findings as PMI increases. Other contributors, including the macromolecular fraction, iron handling, and B0/B1 inhomogeneity, may also affect specific parameters. Overall, our data support a parsimonious model: early water–membrane changes dominate T1; T2 responds more modestly and varies by tissue; T2* is governed by a balance of susceptibility sources and field inhomogeneity, especially in WM ^16,17,36,39,40,42–45^.

Because systematic *ex vivo* PMI studies are not feasible in humans, we used porcine brains as a model, given their cerebral size and gyral complexity ^46^. Although porcine data offer useful insights for human MRI ^47^, age at death (∼6 months) and mode of euthanasia (electric shock, exsanguination) may limit generalizability. Pigs are typically slaughtered via neck incision, which may cause CSF loss. Although the 3D-printed fake “skull” and PBS immersion partially simulated in-situ conditions, rapid postmortem cooling and handling may still differ from human cases ^48^. To facilitate comparison of our porcine results with existing human datasets, we summarized reported qMRI values under various conditions at 7 T (Table 4). T1 values measured in our unfixed *ex vivo* porcine brains are comparable to those published from *in vivo* human MRI (Table 4). oxygenation, and perimortem factors. T1 values measured in our unfixed *ex vivo* porcine brains were comparable to those reported for *in vivo* human MRI ^49^. By contrast, T2 and T2* were longer in unfixed *ex vivo* (porcine) samples than *in vivo* (human) samples ^50,51^. Because there is no standardized preparation pipeline for *ex vivo* MRI after fixation, comparable post-fixation datasets are scarce. Still, relevant information is available from a recent 7 T study of fixed human brains, which employed a different fixation buffer and a different PBS wash duration ^3^. T1 values are comparable with the present data from *ex vivo* porcine brains, whereas T2 and T2* values are longer in the pigs—mirroring the data from unfixed comparison. Overall, T1 values were broadly comparable across species and preparation states, whereas T2 and T2* were consistently longer in *ex vivo* porcine tissue than in human comparators. Nevertheless, the quantitative values are broadly comparable in magnitude.

**Table 4.** GM/WM T1, T2, and T2* at 7 T in porcine *ex vivo* brain vs human literature.

| Condition | Relaxation | Estimated (ms) |  | Literature (ms) |  |
| --- | --- | --- | --- | --- | --- |
|  |  | GM | WM | GM | WM |
| **Unfixed**<br>**or *In Vivo*** | T1 | 1655 ± 46 | 1292 ± 47 | 1686 ± 64<sup>49</sup> | 1218 ± 44<sup>49</sup> |
|  | T2 | 91.19 ± 1.13 | 79.48 ± 2.24 | 55.01 ± 2.07<sup>50</sup> | 39.00 ± 4.43<sup>50</sup> |
|  | T2* | 52.07 ± 1.55 | 36.24 ± 1.33 | 33.2 ± 1.3<sup>51</sup> | 26.8 ± 1.2<sup>51</sup> |
| **Fixed** | T1 | 1126 ± 80 | 887 ± 67 | 1157.67<sup>3</sup> | 899.21<sup>3</sup> |
|  | T2 | 84.94 ± 2.52 | 68.06 ± 2.59 | 47.19<sup>3</sup> | 38.95<sup>3</sup> |
|  | T2* | 62.58 ± 1.73 | 37.50 ± 2.33 | 35.27<sup>3</sup> | 24.19<sup>3</sup> |

GM, gray matter; WM, white matter
Superscript numbers indicate reference sources

This study has several limitations. The accuracy of T1 mapping is a concern. Given time constraints— particularly in unfixed samples—we used the VFA method rather than the more accurate IR method. Insufficient spoiling can bias VFA-based T1 estimates by up to ∼30% relative to IR-derived values ^52^. Other limitations include the lack of within-specimen longitudinal assessment. Additional limitations include TSE-related artifacts, which may affect fitting accuracy. The modest sample size further limits generalizability. Future work with larger datasets may resolve suggestive trends into statistically significant effects.

Despite these limitations, the present study provides important information toward the development of standardized *ex vivo* qMRI technology on fixed samples. Tissue fixation—a complex chemical and physical process ^53^—introduces substantial variability, which has limited the comparability and broader application of *ex vivo* qMRI. In this study, we examined the effects of PMI on unfixed and fixed porcine brains, which have neuroanatomy and MRI properties comparable to those of the human brain. We found that PMI differentially modulates relaxation times across tissue types. The PMI effects were most pronounced within the first 24 h postmortem. Therefore, PMI should be minimized, controlled, and reported transparently in line with the study objectives. For example, T1-focused analyses in GM require rigorous PMI control, as both T1 and GM are particularly sensitive to postmortem delay. Even when fixation attenuates group-level PMI differences due to increased variability, the PMI effects were preserved on T1, irrespective of fixation-induced contrast changes. Thus, standardized workflows are crucial for enhancing reproducibility in postmortem brain imaging. Because hypothermia slows progressive postmortem autolys ^54^, maintaining low temperature prior to autopsy is recommended when considering the application of *ex vivo* MRI. In addition, accelerating the fixation process—such as via perfusion fixation ^55^ or ultrasound acceleration ^56^—may mitigate the PMI effects. Taken together, PMI remains a critical determinant of *ex vivo* qMRI. Complementary strategies—such as temperature management and accelerated fixation—should be employed to enhance the reliability and reproducibility of *ex vivo* qMRI datasets.

## 5. Conclusion

*Ex vivo* qMRI is highly informative; however, results are sensitive to numerous factors, including PMI. PMI effects peak within 24 h postmortem and are strongest for T1 in GM. These effects are attenuated by fixation, which, however, may also amplify inter-sample variability. For studies prioritizing T1-based contrasts in GM or deep structures, investigators should implement active temperature control and accelerated fixation to enhance measurement consistency. Therefore, PMI should be kept to a minimum, strictly regulated, and clearly documented. When rigorous control is not feasible, analyses should be stratified by the PMI window and should explicitly define sequence-specific PMIs. Research should also include PMI as a prespecified covariate and perform sensitivity analyses across PMI windows.

## Acknowledgments

We thank Dr. Tobias Kober (Siemens Healthineers, Erlangen, Germany) for providing the MP2RAGE research prototype sequence used for the 7T scan. Support from the Japan-China Sasakawa Medical Fellowship is gratefully acknowledged.

## Data availability

The data and code underlying this article will be shared by the corresponding author upon reasonable request.

